# Identification of ABHD6 as a regulator of lysophosphatidylserines in the mammalian liver and kidneys

**DOI:** 10.1101/2024.06.02.597019

**Authors:** Arnab Chakraborty, Prajwal Punnamraju, Theja Sajeevan, Arshdeep Kaur, Ullas Kolthur-Seetharam, Siddhesh S. Kamat

## Abstract

Lysophosphatidylserine (lyso-PS) is a potent hormone-like signaling lysophospholipid, which regulates many facets of mammalian biology and dysregulation in its metabolism is associated with several human neurological and autoimmune diseases. Despite the physiological importance and causal relation with human pathophysiology, little is known about the metabolism of lyso-PS in tissues other than the nervous and immune systems. To address this problem, here, we attempted to identify one (or more) lipase(s) capable of degrading lyso-PS in different mammalian tissues. We found that the membrane proteomic fraction of most mammalian tissues possess lyso-PS lipase activity, yet interestingly, the only bona fide lyso-PS lipase ABHD12 displays this enzymatic activity and has control over lyso-PS metabolism only in the mammalian brain. Using an *in vitro* inhibitor screen against membrane proteomic fractions of different tissues, we find that another lipase from the metabolic serine hydrolase family, ABHD6, is a putative lyso-PS lipase in the mouse liver and kidney. Finally, using pharmacological tools, we validate the lyso-PS lipase activity of ABHD6 *in vivo*, and functionally designate this enzyme as a major lyso-PS lipase in primary hepatocytes, and the mammalian liver and kidneys.

**HIGHLIGHTS:**

- Lyso-PS lipase enzymatic activity is present in the membrane proteomic fraction of most mammalian tissues
- An *in vitro* inhibitor screen identifies ABHD6 as a putative lyso-PS lipase in the liver and kidney
- ABHD6 functions as a lyso-PS lipase in primary hepatocytes
- ABHD6 also performs lyso-PS lipase activity *in vivo* and regulates lyso-PS levels in the liver and kidney

## INTRODUCTION

The glycerolysophospholipid lysophosphatidylserine (lyso-PS) is fast emerging as an important signaling lipid, which is central to the functioning of many important biological processes in mammals, including humans^1, 2^. Given its indispensable role in physiology, over the past two decades, the biochemical activities and mechanisms associated with lyso-PS metabolism and signaling have been extensively investigated, mostly in the context of the mammalian central nervous and immune system^1, 2^. This study bias stems from numerous genetic mapping of human subjects (patients) or population-level genome-wide association studies, which causally link the dysregulation in lyso-PS metabolism or signaling to an array of hereditary neurological and autoimmune diseases in humans^1, 3–10^. Despite this, it is now apparent from several recent reports that lyso-PS is quite abundant in almost all mammalian tissues^1^, and yet, very little remains known of its physiological role or metabolism beyond the human nervous and immune systems.

Given its amphiphilic (hydrophilic and lipophilic) chemical nature, lyso-PS acts as a very potent hormone-like signaling molecule^11^. Hence, the physiological concentrations of lyso-PS are tightly regulated by dedicated enzymes in all mammalian tissues^1, 2^. For example, lyso-PS is biosynthesized from phosphatidylserine (PS) precursors by the action of PS-specific phospholipases. In the mammalian central nervous and immune systems, the α/β-hydrolase domain containing protein # 16A (ABHD16A)^12, 13^ and PS-specific phospholipase A1 (PS-PLA1)^14, 15^ have been functionally characterized as PS lipases that tightly control concentrations of intracellular and secreted lyso-PS respectively. On the other hand, lyso-PS signaling in mammals is presumably terminated in all tissues either by its hydrolysis to free fatty acid and glycerophosphoserine by the action of lyso-PS lipases (**Figure 1A**) or via its conversion back to PS by the enzymatic action of lyso-PS specific acyltransferases^1^. To date, in mammals (including humans), the α/β-hydrolase domain containing protein # 12 (ABHD12) remains the only functionally characterized lyso-PS lipase and its functions have been extensively investigated in diverse neurological and immunological settings^11–13, 16–18^. More recently, the lysophosphatidylcholine-specific acyltransferase 3 (LPCAT3) was shown to have selectivity towards arachidonoyl-lyso-PS (C20:4 lyso-PS) in the mammalian brain, where it specifically converts this lyso-PS species to C20:4-containing PS lipids in this tissue^19–21^. Together, these studies show that it is the dynamic interplay between these lyso-PS metabolic enzymes, namely ABHD16A/PS-PLA1, ABHD12, and LPCAT3, that tightly regulates lyso-PS concentrations in the mammalian central nervous and immune systems. While the specifics of mammalian lyso-PS metabolism (both biosynthesis and degradation) have been well worked out in different neurological and immunological paradigms, comparatively, little is known of the enzymes that regulate this metabolism in other peripheral tissues.

**Figure 1.**
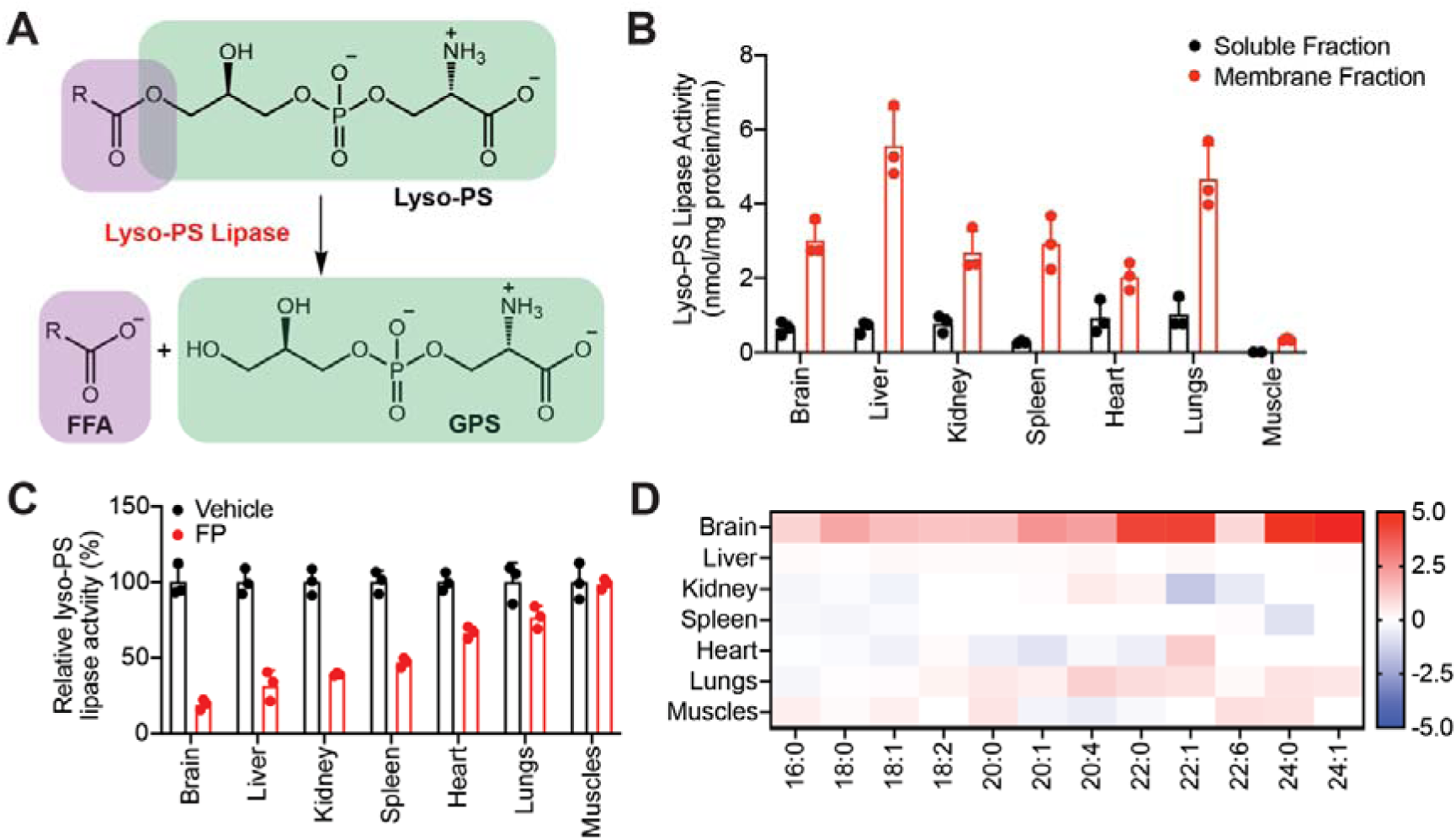
Profiling lyso-PS activity in different mammalian tissues. (**A**) The enzymatic reaction for the conversion of lyso-PS to free fatty acid (FFA) and glycerophosphoserine (GPS) by a lyso-PS lipase. (**B**) The lyso-PS lipase activity in the soluble and membrane proteomic fractions of different mouse tissues. (**C**) The relative lyso-PS lipase activity in the membrane proteomic fractions of different mouse tissues treated with vehicle (DMSO) or FP-rhodamine (20 μM, 45 min). (**D**) Heat map plot showing relative levels of various detectable lyso-PS species from different mouse tissues following deletion of ABHD12. The relative concentration data for each lyso-PS species was obtained by taking the ratio of the knockout value (from ABHD12 knockout mice) to the wild type value (from wild-type mice), where the respective value was calculated as an average from four biological replicates per genotype. The scale adjoining the heat map plot is log_2_[ratio of knockout value to wild type value]. All assays shown in (**B**) and (**C**) were done using 20 μg of proteome against 100 μM C17:1 lyso-PS for 30 min at 37 °C. The bar data presented in (**B**) and (**C**) represents mean ± standard deviation from three biological replicates.

To bridge this knowledge gap, we decided to determine whether peripheral tissues (tissues other than the brain) in mammals have lyso-PS lipase activity (**Figure 1A**), and if ABHD12 and/or any other lipases regulate tissue lyso-PS levels via this enzymatic activity. Using tissue proteomic fractionation and substrate profiling assays, we find that all mouse tissues except the muscle possess lyso-PS lipase activity enriched in the membrane proteomic fraction, and this activity is largely contributed by one (or more) lipase(s) from the metabolic serine hydrolase (mSH) family of enzymes^22, 23^. Further, using ABHD12 knockout mice^18^, we find that this lyso-PS lipase can regulate lyso-PS concentrations only in the brain, and no other tissue. Using a focused inhibitor screen, we find that another lipase from the mSH enzyme family, the α/β-hydrolase domain containing protein # 6 (ABHD6) is possibly a lyso-PS lipase in the mouse liver and kidneys. By pharmacologically inhibiting ABHD6, we show that this lipase controls concentrations of lyso-PS *in vivo* in primary hepatocytes and the liver and kidneys of mice, and thus functionally annotate ABHD6 as a major lyso-PS lipase in these mammalian tissues.

## RESULTS

### Survey of lyso-PS lipase activity in peripheral tissues

While it is now clear that lyso-PS is present ubiquitously in almost all tissues in mammals, most studies pertaining to the metabolism (biosynthesis and degradation) of lyso-PS have surprisingly been restricted to the central nervous and immune systems^1^. Specifically, ABHD12 remains the only bona fide lyso-PS lipase to date^11–13, 16, 18^, and it remains unknown if it can regulate levels of lyso-PSs in different peripheral tissues or if there exist other tissue-resident lipases that can independently regulate the levels of this signaling lysophospholipid in specific tissues. To investigate this, we harvested different mouse tissues (brain, liver, kidney, spleen, heart, lungs, muscle), generated protein lysates, and fractioned these lysates into soluble or membrane proteomic fractions using previously reported protocols^12, 16^. Next, we profiled these different soluble and membrane proteomic fractions using a well-established liquid-chromatography coupled to mass spectrometry (LC-MS) assay to quantify lyso-PS lipase activity in different mouse tissues^11, 16^ (**Figure 1A**). From this profiling study, we found that all tissues except muscle had significant specific lyso-PS lipase activity, that was highly enriched in the membrane proteomic fraction (**Figure 1B, Supplementary Table 1**). Interestingly, relative to the brain membrane lyso-PS lipase activity, this enzymatic activity in the liver and lungs was higher, while that in the kidney, spleen, and heart was comparable (**Figure 1B, Supplementary Table 1**). This result suggests that one (or more) membrane-associated lipase(s) might be responsible for the lyso-PS lipase activities observed in different mouse tissues.

In mammals, the mSH family of enzymes consists of a large number of membrane-associated lipases, a substantial fraction of which still lack any functional assignment^22, 23^. Fluorophosphonate (FP) compounds serve as excellent active-site covalent inhibitors and in turn, activity-based probes for the mSHs. Hence over the years, they have evolved as versatile chemical tools for fishing out unknown biological activities from this enzyme family^23–26^. To determine if one or more mSHs are indeed involved in the lyso-PS lipase activity observed in various mouse tissues, we treated the different tissue membrane proteomic fractions with a FP-probe (FP-rhodamine, 20 μM, 45 min), and subsequently assayed these lysates for lyso-PS lipase activity. From this experiment, we found that, except for the lungs and muscles, the lyso-PS lipase activity in most tissues was inhibited by a FP-probe (**Figure 1C, Supplementary Table 1**). Specifically, FP-probe treatment showed the highest inhibition in the brain, but there was also >50% inhibition of the lyso-PS lipase activity in the membrane proteomic fraction of the liver and kidney (**Figure 1C, Supplementary Table 1**). We also performed the same experiment in the soluble proteomic fractions of different tissues and found that FP-probe treatment did not have any effect (**Supplementary Figure 1A**). Together, this result shows that there is one (or more) membrane-associated mSH(s) that functions as a lyso-PS lipase in different mammalian tissues.

As mentioned previously, studies have now conclusively shown that majority of the brain lyso-PS lipase activity comes from ABHD12, an integral membrane mSH enzyme. Since ABHD12 is the only functionally characterized lyso-PS lipase, we wanted to see if ABHD12 regulates lyso-PS levels in other tissues. Towards this, we first looked at a publicly available gene expression database (biogps.org)^27, 28^ (**Supplementary Figure 1B**), tissue-wide chemoproteomics datasets of mSHs^29^ (**Supplementary Figure 1C**), and performed immunoblotting analysis on membrane proteomic fractions of different mouse tissues described earlier (**Supplementary Figure 1D**). Based on this data, we found that ABHD12 had the highest expression and enzymatic activity in the mammalian brain, and not much in other peripheral tissues. Additionally, we also performed lyso-PS lipase activity assays on the membrane proteomic fractions of different tissues obtained from wild-type or ABHD12 knockout mice, and found that except for the brain, the lyso-PS lipase activities were comparable in all other tissues for both the genotypes (**Supplementary Figure 1E, Supplementary Table 1**). Finally, we measured the lyso-PS levels in different tissues from wild-type and ABHD12 knockout mice and found that except for the brain (where there is a massive accumulation of lyso-PS, particularly very-long chain lyso-PS^11, 12, 18^), ABHD12 deletion did not affect the lyso-PS levels in any other tissue profiled (**Figure 1D, Supplementary Table 1**). Taken together, this suggests that at a tissue-level, ABHD12 regulates the levels of lyso-PS only in the mammalian brain, and that there must exist other lipases from the mSH family which are capable of degrading (hydrolyzing) lyso-PS, and in turn, regulating lyso-PS levels in tissues that possess FP-sensitive lyso-PS lipase activity (e.g. liver, kidney).

### Identification of ABHD6 as a putative lyso-PS lipase

In an effort to identify one (or more) enzyme(s) from different peripheral mouse tissues capable of hydrolyzing lyso-PS, we first collated a focused library of 30 compounds that serve as potent inhibitors (mostly covalent, ranging from very specific to broad spectrum) to lipases from the mSH family^30^ (**Supplementary Table 2**). Next, we treated the membrane proteomic fraction of different mouse tissues that showed ABHD12-independent lyso-PS lipase activity (liver, kidney, spleen, heart and lungs) (**Supplementary Figure 1E, Supplementary Table 1**) with this inhibitor library (10 μM, 45 min), and subsequently assayed them for lyso-PS lipase activity. In this experiment, we considered an inhibitor as a “hit”, if it inhibited the membrane proteomic fraction lyso-PS lipase activity of a particular tissue by >80%. Here, corroborating the results from the lyso-PS lipase assays performed using a FP-probe (**Figure 1C**), we found several hits from the inhibitor library in the membrane proteomic fraction of the liver (4 hits), kidney (4 hits), and spleen (2 hits), but not in the heart and lungs (**Figure 2A**). Since the heart and lungs did not show any hits from the inhibitor library (**Figure 2A**) and are not sensitive to FP-probe treatment (**Figure 1C**), it suggests that the lyso-PS lipase activity in these tissues likely comes from a lipase that is not a mSH enzyme.

**Figure 2.**
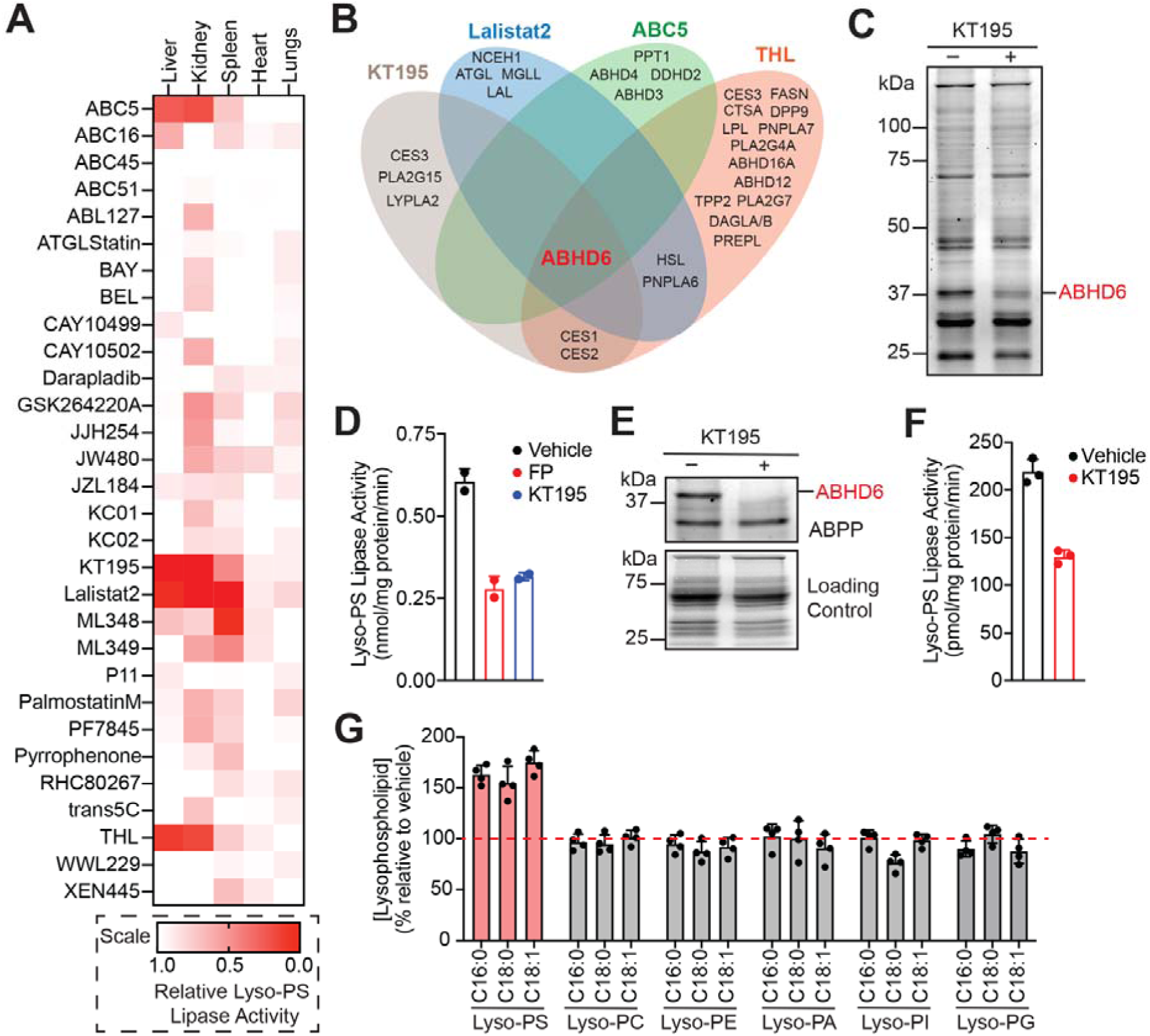
Identification of ABHD6 as a putative lyso-PS lipase. (**A**) Heat map plot showing relative lyso-PS lipase activity from the membrane proteomic fractions of different mouse tissues treated with 30 different lipase inhibitors. The relative lyso-PS lipase activity was obtained by taking the ratio of inhibitor treated samples to vehicle controls, where the respective value was calculated as an average from three biological replicates per group. (**B**) Venn diagram analysis of the various known biological targets of the “hits” obtained from the inhibitor library, showing that ABHD6 is the only common mSH target. (**C**) A representative ABPP gel showing the selective loss of ABHD6 activity in the membrane proteomic fraction in HEK293T cells upon KT195 treatment (1 μM, 4 hours). (**D**) The loss of lyso-PS lipase activity in the membrane proteomic fraction in HEK293T cells upon KT195 treatment (1 μM, 4 hours) from the selective inhibition of ABHD6. FP-rhodamine treated membrane proteomic fraction (*in vitro* treatment, 10 μM, 30 min) was used as a low control in these assay. (**E**) A representative ABPP gel showing the selective loss of ABHD6 activity in the membrane proteomic fraction in primary hepatocytes upon KT195 treatment (1 μM, 4 hours). (**F**) The loss of lyso-PS lipase activity in the membrane proteomic fraction from primary hepatocytes upon KT195 treatment (1 μM, 4 hours) from the selective inhibition of ABHD6. (**G**) Relative levels of different lysophospholipids in primary hepatocytes upon KT195 treatment (1 μM, 4 hours), showing an increase of only lyso-PS, but no other lysophospholipid, from the selective inhibition of ABHD6. The gel-based ABPP experiments shown in (**C**) and (**E**) were performed three times (biological replicates) with reproducible results each time. All assays shown in (**D**) and (**F**) were done using 20 μg of proteome against 100 μM C17:1 lyso-PS for 30 min at 37 °C. The bar data presented in (**D**), (**F**) and (**G**) represents mean ± standard deviation from two (for **D**) or three (**F** and **G**) biological replicates.

Since the liver and kidney showed significant lyso-PS lipase activity that was significantly inhibited by both a FP-probe and compounds from the inhibitor library, we decided to pursue these tissues further. Interestingly, we found that the 4 hits from the inhibitor library in both these tissues were identical (**Figure 2A**), suggesting that, most likely the same mSH regulates the lyso-PS lipase activity in the liver and kidneys. All the 4 hits from the inhibitor library were covalent inhibitors of various mSHs: 2 broad spectrum inhibitors (THL^30, 31^ and Lalistat2^32–34^), and 2 specific inhibitors (KT195 for ABHD6^35, 36^; ABC5 for ABHD4, PPT1^37^), and we decided to assess the overlapping targets between them (**Figure 2B**). Based on publicly available reports and chemoproteomics datasets assessing the targets of these 4 inhibitors, quite surprisingly, we found that the only common mSH target for all these 4 hits was the integral membrane lipase ABHD6 (**Figure 2B**).

Previous studies have shown that recombinant mammalian ABHD6 functions both as a monoacylglycerol lipase^38, 39^ and a lysophospholipase^40^, and we wanted to see if endogenous ABHD6 had any lyso-PS lipase activity. For this, we needed a cell line that was devoid of ABHD12 but had decent ABHD6 expression and activity, and hence chose the HEK293T cell line for this purpose. Treatment of HEK293T cells with the ABHD6-specific inhibitor KT195 (from the inhibitor library) (1 μM, 4 hours) showed the selective loss of ABHD6 by gel-based activity-based protein profiling (ABPP) experiments (**Figure 2C**), and significantly decreased lyso-PS lipase activity (**Figure 2D**) in the membrane proteomic fraction. The loss of lyso-PS lipase activity in HEK293T cells upon KT195 treatment was comparable to the *in vitro* FP-probe treatment (2 μM, 45 min) of the vehicle treated cells, further confirming that all the mSH-associated lyso-PS lipase activity in the membrane proteomic fraction of HEK293T cells was largely coming from ABHD6 (**Figure 2D, Supplementary Table 1**).

Having established that endogenous ABHD6 indeed performs lyso-PS lipase activity, next, we wanted to test a more physiologically relevant cell line to assess if ABHD6 can regulate lyso-PS levels. Since ABHD12 has negligible activity in the liver (**Supplementary Figure 1C**), we chose to study the role of ABHD6 in modulating lyso-PS lipase activity and lyso-PS levels in primary hepatocytes. Treatment of primary hepatocytes with KT195 (1 μM, 4 hours) showed complete inhibition of ABHD6 in the membrane proteomic fraction by gel based ABPP (**Figure 2E**). Concomitant to the inhibition of ABHD6, we found that the lyso-PS lipase activity associated with the membrane proteomic fraction of primary hepatocytes also substantially decreased upon the same KT195 treatment (**Figure 2F, Supplementary Table 1**). Since previous studies have shown that ABHD6 functions as a general lysophospholipase^40^, besides lyso-PS, we also decided to measure the levels of other lysophospholipids (lyso-PC, lyso-PE, lyso-PA, lyso-PI, and lyso-PG) in primary hepatocytes upon KT195 treatment. Interestingly, here we found that relative to the vehicle control, KT195 treated primary hepatocytes resulted in the accumulation of only lyso-PS (> 1.5-fold), but no other lysophospholipid (**Figure 2G**). Taken together, these studies suggest that ABHD6 functions as putative lyso-PS lipase in primary hepatocytes, and selectively regulates the levels of only this lysophospholipid in these liver cells.

### ABHD6 regulates lyso-PS levels in the liver and kidney

A large-scale gene expression database (biogps.org)^27, 28^ (**Supplementary Figure 2A**) and ABPP analysis of mSHs^29^ (**Supplementary Figure 2B**) in different mouse tissues have shown that ABHD6 has the highest activity in the membrane proteomic fraction of the brain and liver, and to a lesser extent in the kidneys. Hence, having demonstrated ABHD6’s role in the regulation of lyso-PS levels in primary hepatocytes, next, we wanted to see if it performs lyso-PS lipase activity *in vivo* (in the liver and maybe other tissues). Since it has been reported that KT195 is poorly bioavailable in mice models, we chose a structurally related *in vivo* active ABHD6-specific inhibitor KT185^36^, to assess the effect of pharmacological inhibition of ABHD6 on lyso-PS metabolism in different mouse tissues. In the experiment, upon orally dosing mice with KT185 (10 mg/kg body weight, 4 hours), we harvested different tissues and assessed them *ex vivo*.

To determine the efficacy of KT185 *in vivo*, using gel based ABPP, we checked for the inhibition of ABHD6 in the membrane proteomic fractions of different mouse tissues (**Figure 3A**). Consistent with available expression and enzyme activity data, we were able to reliably detect ABHD6 activity in the membrane proteomic fractions of the brain, liver and kidney, and found that KT185 treatment (10 mg/kg body weight, 4 hours) in mice resulted in the complete inhibition of ABHD6 in these tissues (**Figure 3A**). However, we were unable to detect any ABHD6 activity in the membrane proteomic fractions of spleen, heart or lungs in the same gel-based ABPP experiments (**Figure 3A**). Concomitant to the inhibition of ABHD6 in the liver and kidney, next, we found that the lyso-PS lipase activity in the membrane proteomic fractions of these tissues was also substantially reduced (∼ 50% and 40% in the liver and kidneys respectively) upon KT185 treatment in mice (**Figure 3B, Supplementary Table 1**). Like the gel-based ABPP assays, we found no changes in the lyso-PS lipase activity in the spleen, heart or lungs upon KT185 treatment in mice (**Figure 3B, Supplementary Table 1**). Interestingly, we found that despite the complete inhibition of ABHD6 by KT185 treatment, there was no change in the lyso-PS lipase activity in the brain membrane proteomic fraction, further confirming that ABHD12 is the major lyso-PS lipase in the mammalian brain^11–13, 18^, and perhaps ABHD6 has another biochemical function in the central nervous system^41^ (**Figure 3B, Supplementary Table 1**).

**Figure 3.**
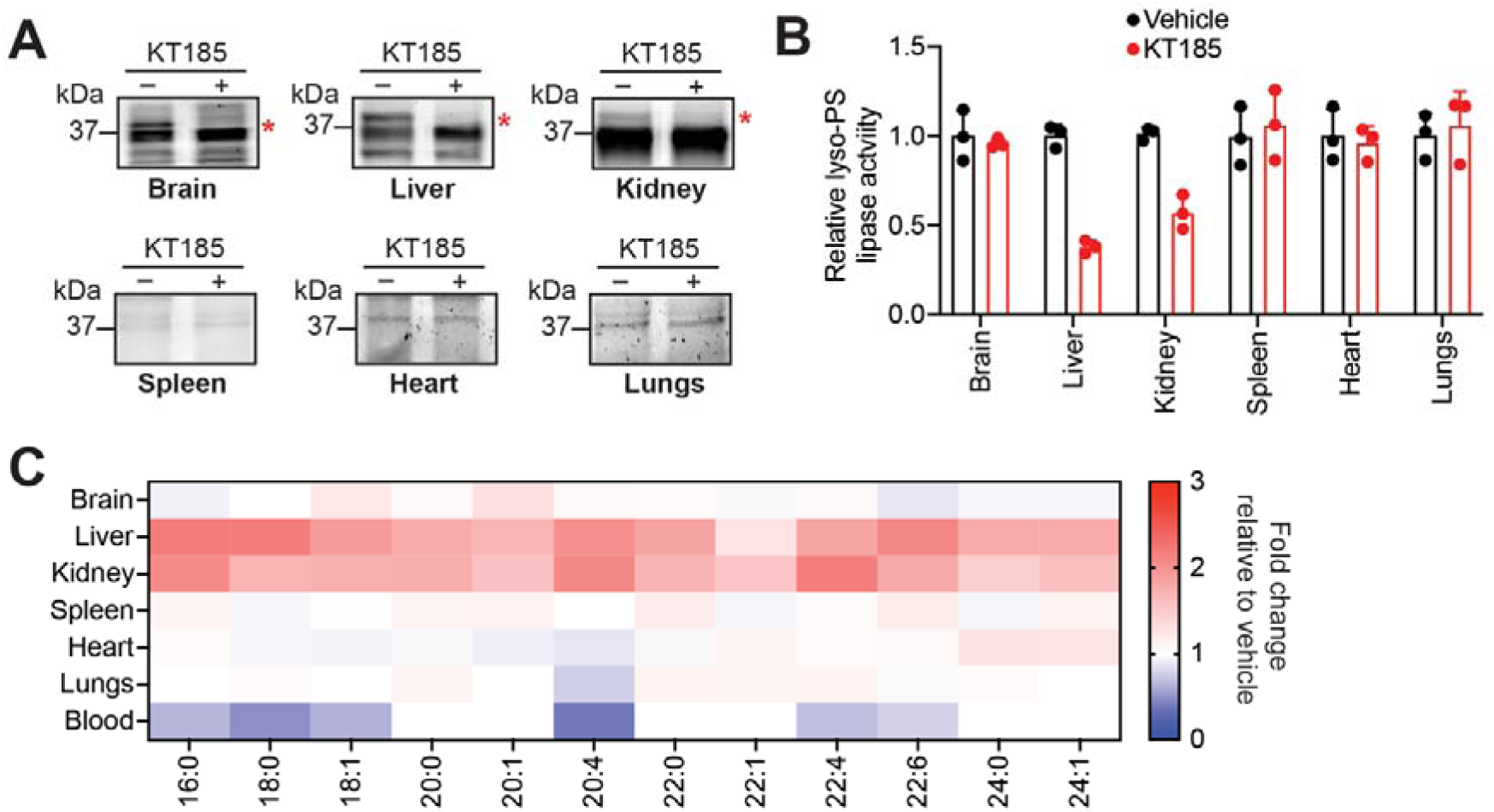
ABHD6 functions as a lyso-PS lipase in the liver and kidney. (**A**) Representative ABPP gels (zoomed) showing the selective loss of ABHD6 activity in the membrane proteomic fraction of the brain, liver and kidney upon KT185 treatment (10 mg/kg body weight, 4 hours). These gel-based ABPP experiments were performed three times (biological replicates) with reproducible results each time. (**B**) The relative lyso-PS lipase activity in the membrane proteomic fractions of different mouse tissues treated with vehicle or KT185 (10 mg/kg body weight, 4 hours) following ABHD6 inhibition. All assays were done using 20 μg of proteome against 100 μM C17:1 lyso-PS for 30 min at 37 °C. The bar data represents mean ± standard deviation from three biological replicates. (**C**) Heat map plot showing relative levels of various detectable lyso-PS species from different mouse tissues following inhibition of ABHD6 by KT185 treatment (10 mg/kg body weight, 4 hours). The relative concentration data for each lyso-PS species was obtained by taking the ratio of the KT185 treated value (from ABHD6 inhibition) to the vehicle treated value, where the respective value was calculated as an average from at least four biological replicates per genotype. The scale adjoining the heat map plot is [ratio of KT185 treated value to vehicle treated value].

Finally, we measured the lyso-PS concentrations in different tissues following KT185 treatment in mice. Here, consistent with complete ABHD6 inhibition and the loss of lyso-PS lipase activity in the liver and kidney, relative to the vehicle control, we found that both these tissues showed a significant accumulation (∼ 2 to 3-fold) of different lyso-PS species (**Figure 3C, Supplementary Table 1**). Corroborating the gel-based ABPP assays and the lyso-PS lipase activity profiles, we found that all other tissues (brain, spleen, heart and lungs) showed little to no changes in tissue concentrations of lyso-PS (**Figure 3C, Supplementary Table 1**). Further, in the liver and kidneys, we also measured the concentrations of several other lysophospholipids, and found that these showed negligible changes upon KT185 treatment in mice (**Supplementary Figure 2C, 2D**). Quite surprisingly, we found that KT185 treatment in mice resulted in significantly decreased levels of lyso-PS circulating in the blood (**Figure 3C, Supplementary Table 1**), suggesting that ABHD6 activity perhaps has a role to play in the secretion of lyso-PS lipids from the liver and kidneys. Taken together, our gel-based ABPP assays, lyso-PS lipase activity profiles, and lyso-PS measurements in different tissues following KT185 treatment in mice strongly suggest that *in vivo* ABHD6 functions as a major lyso-PS lipase in the liver and kidneys.

## DISCUSSION

Signaling lipids (e.g.: lysophospholipids) are indispensable systemic small-molecule mediators of essential biological processes in mammals, and dysregulation in their metabolism and/or signaling has clearly established links to an array of human diseases^42–44^. Over the past two decades, lyso-PS has emerged as yet another biomedically important signaling lysophospholipid, and most of the knowledge pertaining to its metabolism and signaling is restricted to the central nervous and immune systems, given its involvement in several human neurological and autoimmune disorders^1^. Specifically, in the central nervous system, lyso-PS controls the activation of microglial cells, and in turn neuroinflammation, particularly in the cerebellum. On the other hand, in the immune system, this signaling lysophospholipid regulates histamine release from mast cell degranulation, pro-inflammatory responses from macrophages, and the maturation of T cells, to list a few processes^1, 2^. While it is now evident that lyso-PS is ubiquitously present^1^, little remains known on how it is enzymatically produced and metabolized, or what are its physiological functions in other (peripheral) mammalian tissues.

To address this problem in part, we decided to investigate the degradation of this signaling lipid via a lyso-PS lipase activity assay in different mammalian tissues (**Figure 1A**). We found that other than the muscles, all other mammalian tissues possess significant lyso-PS lipase activity that is enriched in the membrane proteomic fraction (**Figure 1B**). Further profiling suggested that one (or more) mSH enzyme(s) was largely responsible for this activity in the brain, liver, kidney and spleen (**Figure 1C**). Using a combination of biochemical assays and tissue lyso-PS measurements coupled with protein expression data, we show that the only annotated lyso-PS lipase ABHD12 controls lyso-PS levels only in the brain (and central nervous system^12, 18^), and no other peripheral tissue (**Figure 1D, Supplementary Figure 1**). In a quest to identify lipases in peripheral tissues capable of hydrolyzing lyso-PS, we screened the membrane proteomic lysates of different tissues against a focused library of inhibitors of lipases belonging to the mSH family (**Figure 2A**). From this screen, we found that the integral membrane mSH enzyme ABHD6 was a potential lyso-PS lipase in the liver and kidneys (**Figure 2B**). Following this, we validated the ability of ABHD6 to regulate lyso-PS levels in mammalian cells, particularly in primary hepatocytes, where we found that via its lysophospholipase activity, ABHD6 selectively controls levels of only lyso-PS and no other lysophospholipid (**Figure 2C-G**). Lastly, using a selective inhibitor in mice, we show that the loss of ABHD6 activity results in significantly decreased lyso-PS lipase activity, and a concomitant increase in lyso-PS levels, but no other lysophospholipid, only in the liver and kidney (**Figure 3**). Taken together, we provide strong biochemical evidence in support of ABHD6’s ability to function selectively as a lyso-PS lipase in the liver and kidney.

Moving ahead, our studies open several new research opportunities. For instance, in the central nervous system, ABHD6 is designated as a key lipase involved in the metabolism of the endocannabinoid 2-arachidonoylglycerol (2-AG)^38, 41, 45^. However, it has shown that the long-term loss of ABHD6 (via knock-down) does not alter 2-AG levels in peripheral tissues (e.g. liver, kidney), but results in substantially altered global lipid profiles in the liver, especially under conditions of a high-fat diet^40, 46^. Further, ABHD6 is tentatively annotated as a promiscuous lysophospholipase, and shown to function as a general regulator of glycerophospholipid metabolism in the liver in mice models recapitulating high-fat diet induced obesity^40^. Our studies partly corroborate these findings, and show that ABHD6 functions as a lyso-PS-specific lysophospholipase in the liver and kidney in mice, and selectively controls levels of lyso-PS lipids, but not other lysophospholipids, in these tissues under normal physiological conditions (not in high-fat diet paradigms) (**Figure 3**, **Supplementary Figure 2**). Given its recent links to systemic (glucose) metabolism^1^, it will be interesting to see how dysregulated lyso-PS signaling in the liver might contribute towards metabolic conditions such as hepatic steatosis or systemic insulin resistance (modulated by ABHD6 activity), and crosstalk with other lipid pathways involved in such human diseases.

Our profiling studies from different mammalian tissues also show that despite having abundant lyso-PS levels, the muscles do not possess any measurable lyso-PS lipase activity (**Figure 1B**). This result suggests that in muscles, lyso-PS may likely be metabolized via an acyltransferase-type activity. The identification of such a lysophospholipid acyltransferase will shed new insights into lyso-PS metabolism in tissues, where lyso-PS lipases are unable to regulate its physiological concentrations. Recent studies have annotated the lysophospholipid acyltransferase LPCAT3 as an enzyme capable of metabolizing lyso-PS, and shown a functional crosstalk between LPCAT3 and ABHD12 in the mammalian brain^21^. Of note, publicly available gene expression databases show that LPCAT3 has abundant expression in the kidney, liver, brain and skeletal muscles^27, 28^. Further, recent studies in mice show an important role for this enzyme in regulating hepatic lipid secretion^19, 20, 47, 48^. Hence, it will be interesting to see if LPCAT3 or some other lyso-PS specific acyltransferase can metabolize lyso-PS and regulate its physiological concentrations in the muscles. Additionally, given its expression profiles, and similar to studies done with ABHD12 in the brain^21^, it will be important to understand the functional interplay, if any, between LPCAT3 and ABHD6 in regulating lyso-PS metabolism and signaling in the liver and kidney.

Finally, our studies show that the lyso-PS lipase activity from the heart and lungs is not sensitive to a FP-probe (**Figure 1C**) or selected inhibitors from the screen (**Figure 2A**). This suggests that the putative lipase(s) is most likely not a member of the mSH family, and thus opens new directions towards finding the identity of this enzyme(s). We also find from the inhibitor screen, that the lyso-PS lipase activity from the membrane proteomic lysate of the spleen is almost completely inhibited by ML348 (**Figure 2A**), a potent yet selective inhibitor of lysophospholipase 1 (LYPLA1) enzyme^49, 50^. Since LYPLA1 has ubiquitous expression in all tissues and is poorly characterized in terms of its lysophospholipase activity^51, 52^, it will also be worthwhile determining if this lipase indeed functions as a lyso-PS in the spleen, and if so, how it crosstalks with the immune system. Taken together, given the biomedical importance of lyso-PSs, and their direct association to human diseases, our studies illuminate new avenues of research in understanding the metabolism and signaling pathways regulated by this emerging lysophospholipid beyond the mammalian central nervous and immune systems.

## SIGNIFICANCE

Lyso-PS is a hormone-like signaling lysophospholipid that controls many facets of mammalian biology. Given its association with numerous neurological and autoimmune disorders in humans, most studies pertaining to the metabolism of lyso-PS and its signaling pathways remain restricted to the mammalian central nervous and immune systems, despite the ubiquitous presence of lyso-PS in all mammalian tissues. In a pursuit to understand the enzymatic degradation of lyso-PS, we profiled the lyso-PS lipase activity associated with different mammalian tissues. We show that besides muscles, all other tissues possess significant lyso-PS lipase activity, largely coming from membrane-associated mSH enzymes. We show conclusively that via its enzymatic activity, the only functionally characterized lyso-PS lipase ABHD12 controls lyso-PS levels only in the mammalian brain, and no other tissue. Using a focused inhibitor screen in conjunction with follow up pharmacological studies in primary hepatocytes and mice, we identify ABHD6 as a major lyso-PS lipase in the liver and kidney, where it selectively regulates concentrations of lyso-PS, and no other lysophospholipid. Our findings thus show how lyso-PS is metabolized in peripheral tissues, and open new avenues for studying this emerging signaling lysophospholipid in the context of systemic metabolism and metabolic diseases in humans.

## MATERIALS AND METHODS

### Materials

Unless otherwise mentioned, all chemicals, buffers, and reagents were procured from Sigma Aldrich; all lipid standards were purchased from Avanti Polar Lipids; all mammalian tissue culture media and consumables were purchased from HiMedia; and all LC-MS grade solvents were purchased from JT Baker. Where ever applicable, we mention the catalog numbers of the materials used in the respective section below.

### Mice Experiments

All mouse studies and experiments described in this paper have received formal approval from the Indian Institute of Science Education and Research, Pune – Institutional Animal Ethics Committee (application nos: IISER_Pune IAEC/2021_01/09; IISER_Pune IAEC/202_01/05) constituted as per the guidelines and norms provided by the Committee for the Purpose of Control and Supervision of Experiments in Animals, Government of India. All experimental mice were housed in the National Facility for Gene Function in Health and Disease (NFGFHD), IISER Pune, and were studied between 2 – 3 months of age. All mice used in this study were from the C57BL/6J genetic background (RRID: IMSR_JAX:000664), and had *ab libitum* access to food and water. In all experiments, equal number of male and female mice were used. For studies involving ABHD12 knockout mice, age/sex matched littermates were used as controls. Primary hepatocytes were generated using established protocols previously reported by us^53–55^.

### Preparation of Tissue Proteomic Fractions

Mice were deeply anesthetized with isoflurane, and euthanized by cervical dislocation. Thereafter, the tissues of interest were surgically harvested, washed with cold sterile Dulbecco’s phosphate buffered saline (DPBS) (HiMedia, catalog no: TL1006) (three times), weighed, flash-frozen in liquid nitrogen, and stored at −80 °C till further use. The pre-weighed tissues were thawed on ice, resuspended in 500 μL of ice-cold DPBS, and homogenized using a Bullet Blender 24 (Next Advance) using one scoop of 0.5-mm diameter glass beads (Next Advance) for brains, one scoop of 2.3-mm diameter stainless steel beads (Next Advance) for the liver, heart and spleen, and one scoop of 1-mm diameter zirconium oxide beads (Next Advance) for the kidneys and lungs, at a speed setting of 8 for 3 min at 4 °C (two times). To this tissue homogenate, 500 μL of cold sterile DPBS was added and mixed by pipetting. These homogenates were subsequently probe sonicated for 90 seconds (2 seconds on/off pulses at 60% amplitude) using a medium-sized probe, and these lysates were centrifuged at 1000g for 5 minutes at 4 °C to separate the beads and tissue debris from the tissue proteome. The resulting tissue lysate (supernatant, ∼800 μL) was separated by pipetting and centrifuged at 100,000g for 45 minutes at 4 °C in an Optima MAX-XP Beckman coulter ultracentrifuge. Following ultracentrifugation, the resulting supernatant (∼500 μL) was separated by pipetting and labeled as the soluble proteomic fraction. The resulting pellet was washed three times with cold sterile DPBS, and resuspended in 400 μL cold sterile DPBS by sonication. The resulting lysate was labeled as the membrane proteomic fraction for a particular tissue. The protein concentrations of the membrane and soluble proteomic fractions were measured using the Pierce BCA Protein Assay Kits (Thermo Fisher Scientific, catalog no: 23225).

### Preparation of Cellular Proteomic Fractions

HEK293T (RRID: CVCL_0063, female) cells were purchased from the ATCC, and cultured and maintained as reported earlier^11^. Primary hepatocytes were generated using established protocols previously reported by us^53–55^. All mammalian cells were harvested by scraping, and washed three-times with cold sterile DPBS. The cellular protein lysates were prepared by resuspending the cells in 500 μL DPBS, and probe sonicating them for 30 seconds (2 seconds on/off pulses at 60% amplitude) using a medium-sized probe. These lysates were subsequently centrifuged at 1000g for 5 minutes at 4 °C to separate the unlysed cells and debris from the proteome. The resulting cell lysate (supernatant, ∼400 μL) was separated by pipetting and centrifuged at 100,000g for 45 minutes at 4 °C in an Optima MAX-XP Beckman coulter ultracentrifuge. Following ultracentrifugation, the resulting supernatant (∼300 μL) was separated by pipetting and labeled as the soluble proteomic fraction. The resulting pellet was washed three times with cold sterile DPBS, and resuspended in 400 μL cold sterile DPBS by sonication. This resulting lysate was labeled as the membrane proteomic fraction for particular mammalian cells. The protein concentrations of the membrane and soluble proteomic fractions were measured using the Pierce BCA Protein Assay Kits (Thermo Fisher Scientific, catalog no: 23225).

### Lyso-PS Lipase Substrate Assays

All lyso-PS lipase substrate assays were done using LC-MS protocols previously reported by us^11, 12, 30^. Briefly, 20 μg of the soluble or membrane proteomic fraction obtained from mammalian tissues or cells was incubated with 100 μM lyso-PS substrate (C17:1 lyso-PS; Avanti Polar Lipids, catalog no: 858141) in DPBS to a final volume of 100 μL at 37 °C with constant shaking (Eppendorf, Thermo Mixer C). After letting the reaction proceed to 30 minutes, it was quenched with 300 μL of 2:1 chloroform (CHCl_3_): methanol (MeOH) spiked with an internal standard (0.5 nmol of C15:0 free fatty acid (FFA); Sigma Aldrich, catalog no: P6125). The mixture was vortexed vigorously and centrifuged at 2300g to separate the aqueous (top) and organic (bottom) phases. The organic phase was dried under a stream of nitrogen gas and resolubilized in 150 μL of 2:1 CHCl_3_: MeOH for LC-MS analysis. This organic extract was injected into an Agilent G6125B single quadrupole LC/MSD and analyzed in the negative ion mode using an electrospray ionization (ESI) source. The LC separation was performed using a Gemini 5U C-18 column (Phenomenex) coupled with a Gemini guard column (Phenomenex, 4×3 mm, Phenomenex security cartridge). The solvents used were buffer A: 95:5 = Water: MeOH; and buffer B: 60:35:5 Isopropanol (IPA): MeOH: water. All runs were 15 minutes, starting with 0.3 mL/min 100% buffer A for 1.5 minutes, 0.5 mL/min linear gradient to 100% buffer B over 5 minutes, 0.5 mL/min 100% buffer B for 5.5 minutes, and equilibration with 0.5 mL/min 100% buffer A for 3 minutes. The following MS parameters were used: Drying gas flow = 10 L/min, nebulizer pressure = 45 Ψ, drying gas temperature = 250 °C, capillary voltage = 4 kV. The product release was quantified by measuring the area under the curve for the peak corresponding to C17:1 FFA (produced from C17:1 lyso-PS), and normalizing it to the internal standard (C15:0 FFA). The substrate hydrolysis rate was corrected by subtracting the non-enzymatic rate of hydrolysis, which was obtained by using heat-denatured (15 min at 95 °C) control proteomes as reported earlier^11, 12, 30^.

### Inhibitor Screen in Tissue Membrane Proteomic Fractions

Membrane proteomic fractions (50 μL of 1 mg/mL) from different mouse tissues were incubated with different lipase inhibitors from the focused library **(Supplementary Table 2)** at a concentration of 10 μM for 45 minutes. Following this, the lysates were subsequently assayed for lyso-PS lipase activity as described earlier. DMSO was used as a vehicle control (high control), whereas FP-rhodamine treatment (10 μM for 45 minutes; in case of liver, kidney, spleen) or heat denatured proteome (in case of heart, lungs) was used as loss of lyso-PS lipase control (low control) in this inhibitor screen.

### Lyso-PS Measurements in Cells and Tissues

Phospholipids, including lyso-PS, were extracted from mammalian cells or tissues using a modified Folch lipid extraction procedure previously reported by us^11, 12, 30^. During the extraction, C17:1 lyso-PS (1 nmol per sample) was used as an internal standard. The dried lipid extract was resuspended in 200 μL 2:1 CHCl_3_: MeOH, and 10 μL of this was injected into an Agilent Technologies 6470 Triple Quadrupole LC/MS for quantification of lyso-PS using established multiple reaction monitoring (MRM) transition-based analysis previously reported by us^11, 12, 30^. The LC parameters and runtime were identical to those described for the lyso-PS lipase substrate hydrolysis assays. All LC-MS analysis was performed in negative ion mode using ESI source the following parameters: drying and sheath gas temperature = 320 °C; drying and sheath gas flow = 10 L/min; nebulizer pressure = 45 Ψ; fragmentor voltage = 160 V; capillary voltage = 3000 V; and nozzle voltage = 1000 V. All lyso-PS species were quantified by normalizing their respective areas under the curve and normalizing it to the area under the curve of the internal standard (C17:1 lyso-PS) and then normalizing to tissue weight or total cellular proteins.

### Untargeted Lipid Measurements

The dried lipid extracts described in the previous section were resolubilized in 200 μL of 2:1 CHCl_3_: MeOH and 10 μL was injected into an Agilent 6545 Quadrupole Time-Of-Flight (QTOF) LC-MS/MS for semi-quantitative analysis using high-resolution auto MS/MS methods and chromatography techniques. A Gemini 5U C-18 column (Phenomenex) coupled with a Gemini guard column (Phenomenex, 4×3 mm, Phenomenex security cartridge) was used for LC separation. The solvents used for the LC-MS analysis were buffer A: 95:5 water: MeOH, and buffer B: 60:35:5 IPA: MeOH: water. For negative ion mode analysis, 0.1% (v/v) ammonium hydroxide was added in each buffer, while 0.1% (v/v) formic acid + 10 mM ammonium formate were used as additives for analysis in the positive ion mode. All untargeted lipid measurement runs were 60 minutes, starting with 0.3 mL/min 100% buffer A for 5 minutes, 0.5 mL/min linear gradient to 100% buffer B over 40 minutes, 0.5 mL/min 100% buffer B for 10 minutes, and then equilibration with 0.5 mL/min 100% buffer A for 5 minutes. All LC-MS runs were performed using ESI source with the following MS parameters: drying and sheath gas temperature = 320 °C; drying and sheath gas flow rate = 10L/min; fragmentor voltage = 150 V; capillary voltage = 4 kV; nebulizer (ion source gas) pressure = 45 Ψ and nozzle voltage = 1 kV. For analysis of different lipids, particularly lysophospholipids, a curated lipid library was employed in the form of a Personal Compound Database Library (PCDL), and the peaks were validated based on relative retention times and MS/MS fragments obtained. All lipid species were quantified by normalizing areas under the curve relative to the internal standard added and then normalized either to the tissue weight or total cellular proteins.

### Gel-based ABPP Assays

All gel-based ABPP assays were performed using protocols previously reported by us^11, 56, 57^. Briefly, 50 μL of 1mg/mL soluble or membrane proteomic fraction from mammalian cells or tissues were incubated with 2 μM FP-rhodamine for 45 minutes at 37°C with constant shaking (Eppendorf, Thermo Mixer C). The reactions were quenched by adding 20 μL of 4x SDS-PAGE loading buffer followed by boiling for 5 minutes, and 40μL of this sample was loaded on a 12.5% SDS-PAGE gel. Competitive gel-based ABPP experiments were done as reported earlier^11, 56, 57^. All gels were visualized using an iBright1500 gel documentation system (Invitrogen).

## ACKNOWLEDGEMENTS

We gratefully acknowledge support from the Science and Engineering Research Board (SERB), Department of Science and Technology, Govt. of India (Grants: SB/SJF/2021-22/01 and CRG/2020/000023 to S.S.K.), an EMBO Young Investigator Award (to S.S.K.), and the Council of Scientific and Industrial Research (CSIR), Govt. of India (Graduate Student Fellowship to A.C.). We thank Saddam Shekh for technical assistance and maintenance of the biological mass spectrometry facility at IISER Pune. All staff members of the National Facility for Gene Function in Health and Disease (NFGFHD) at IISER Pune (supported by a grant from the Department of Biotechnology, Govt. of India; BT/INF/22/SP17358/2016) are thanked for maintaining and providing mice for this study. Members of the S.S.K. lab at IISER Pune are thanked for providing critical comments and inputs throughout the course of this study.

## AUTHOR CONTRIBUTIONS

A.C. performed all the experiments with help from P.P. and T.S.; A.K. and U.K.S. helped generate the primary hepatocytes used in this study. A.C. and S.S.K. conceived the project and wrote the paper with inputs from all authors. S.S.K. supervised and acquired funding for the project.

## REFERENCES

1. Chakraborty, A., and Kamat, S.S. (2024). Lysophosphatidylserine: A Signaling Lipid with Implications in Human Diseases. Chem Rev 124, 5470–5504. 10.1021/acs.chemrev.3c00701.

2. Shanbhag, K., Mhetre, A., Khandelwal, N., and Kamat, S.S. (2020). The Lysophosphatidylserines-An Emerging Class of Signalling Lysophospholipids. J Membr Biol 253, 381–397. 10.1007/s00232-020-00133-2.

3. Fiskerstrand, T., H’Mida-Ben Brahim, D., Johansson, S., M’Zahem, A., Haukanes, B.I., Drouot, N., Zimmermann, J., Cole, A.J., Vedeler, C., Bredrup, C., et al. (2010). Mutations in ABHD12 cause the neurodegenerative disease PHARC: An inborn error of endocannabinoid metabolism. Am J Hum Genet. 87, 410–417.

4. Fiskerstrand, T., Knappskog, P., Majewski, J., Wanders, R.J., Boman, H., and Bindoff, L.A. (2009). A novel Refsum-like disorder that maps to chromosome 20. Neurology 72, 20–27.

5. Miyake, N., Silva, S., Troncoso, M., Okamoto, N., Andachi, Y., Kato, M., Iwabuchi, C., Hirose, M., Fujita, A., Uchiyama, Y., and Matsumoto, N. (2022). A homozygous ABHD16A variant causes a complex hereditary spastic paraplegia with developmental delay, absent speech, and characteristic face. Clin Genet 101, 359–363. 10.1111/cge.14097.

6. Lemire, G., Ito, Y.A., Marshall, A.E., Chrestian, N., Stanley, V., Brady, L., Tarnopolsky, M., Curry, C.J., Hartley, T., Mears, W., et al. (2021). ABHD16A deficiency causes a complicated form of hereditary spastic paraplegia associated with intellectual disability and cerebral anomalies. American journal of human genetics 108, 2017–2023. 10.1016/j.ajhg.2021.09.005.

7. Yahia, A., Elsayed, L.E.O., Valter, R., Hamed, A.A.A., Mohammed, I.N., Elseed, M.A., Salih, M.A., Esteves, T., Auger, N., Abubaker, R., et al. (2021). Pathogenic Variants in ABHD16A Cause a Novel Psychomotor Developmental Disorder With Spastic Paraplegia. Front Neurol 12, 720201. 10.3389/fneur.2021.720201.

8. Zhao, Y., Hasse, S., and Bourgoin, S.G. (2021). Phosphatidylserine-specific phospholipase A1: A friend or the devil in disguise. Prog Lipid Res 83, 101112. 10.1016/j.plipres.2021.101112.

9. Szymanski, K., Miskiewicz, P., Pirko, K., Jurecka-Lubieniecka, B., Kula, D., Hasse-Lazar, K., Krajewski, P., Bednarczuk, T., and Ploski, R. (2014). rs3827440, a nonsynonymous single nucleotide polymorphism within GPR174 gene in X chromosome, is associated with Graves’ disease in Polish Caucasian population. Tissue antigens 83, 41–44. 10.1111/tan.12259.

10. Chu, X., Shen, M., Xie, F., Miao, X.J., Shou, W.H., Liu, L., Yang, P.P., Bai, Y.N., Zhang, K.Y., Yang, L., et al. (2013). An X chromosome-wide association analysis identifies variants in GPR174 as a risk factor for Graves’ disease. Journal of medical genetics 50, 479–485. 10.1136/jmedgenet-2013-101595.

11. Khandelwal, N., Shaikh, M., Mhetre, A., Singh, S., Sajeevan, T., Joshi, A., Balaji, K.N., Chakrapani, H., and Kamat, S.S. (2021). Fatty acid chain length drives lysophosphatidylserine-dependent immunological outputs. Cell Chem Biol 28, 1169–1179. 10.1016/j.chembiol.2021.01.008.

12. Singh, S., Joshi, A., and Kamat, S.S. (2020). Mapping the Neuroanatomy of ABHD16A, ABHD12, and Lysophosphatidylserines Provides New Insights into the Pathophysiology of the Human Neurological Disorder PHARC. Biochemistry 59, 2299–2311. 10.1021/acs.biochem.0c00349.

13. Kamat, S.S., Camara, K., Parsons, W.H., Chen, D.H., Dix, M.M., Bird, T.D., Howell, A.R., and Cravatt, B.F. (2015). Immunomodulatory lysophosphatidylserines are regulated by ABHD16A and ABHD12 interplay. Nat Chem Biol 11, 164–171. 10.1038/nchembio.1721.

14. Aoki, J., Inoue, A., Makide, K., Saiki, N., and Arai, H. (2007). Structure and function of extracellular phospholipase A1 belonging to the pancreatic lipase gene family. Biochimie 89, 197–204. 10.1016/j.biochi.2006.09.021.

15. Nagai, Y., Aoki, J., Sato, T., Amano, K., Matsuda, Y., Arai, H., and Inoue, K. (1999). An alternative splicing form of phosphatidylserine-specific phospholipase A1 that exhibits lysophosphatidylserine-specific lysophospholipase activity in humans. J Biol Chem 274, 11053–11059. 10.1074/jbc.274.16.11053.

16. Joshi, A., Shaikh, M., Singh, S., Rajendran, A., Mhetre, A., and Kamat, S.S. (2018). Biochemical characterization of the PHARC-associated serine hydrolase ABHD12 reveals its preference for very-long-chain lipids. Journal of Biological Chemistry 293, 16953–16963. 10.1074/jbc.RA118.005640.

17. Ogasawara, D., Ichu, T.A., Vartabedian, V.F., Benthuysen, J., Jing, H., Reed, A., Ulanovskaya, O.A., Hulce, J.J., Roberts, A., Brown, S., et al. (2018). Selective blockade of the lyso-PS lipase ABHD12 stimulates immune responses in vivo. Nature chemical biology 14, 1099–1108. 10.1038/s41589-018-0155-8.

18. Blankman, J.L., Long, J.Z., Trauger, S.A., Siuzdak, G., and Cravatt, B.F. (2013). ABHD12 controls brain lysophosphatidylserine pathways that are deregulated in a murine model of the neurodegenerative disease PHARC. Proc Natl Acad Sci U S A 110, 1500–1505. 10.1073/pnas.1217121110.

19. Lagrost, L., and Masson, D. (2022). The expanding role of lyso-phosphatidylcholine acyltransferase-3 (LPCAT3), a phospholipid remodeling enzyme, in health and disease. Curr Opin Lipidol 33, 193–198. 10.1097/MOL.0000000000000820.

20. Shao, G., Qian, Y., Lu, L., Liu, Y., Wu, T., Ji, G., and Xu, H. (2022). Research progress in the role and mechanism of LPCAT3 in metabolic related diseases and cancer. J Cancer 13, 2430–2439. 10.7150/jca.71619.

21. Ichu, T.A., Reed, A., Ogasawara, D., Ulanovskaya, O., Roberts, A., Aguirre, C.A., Bar-Peled, L., Gao, J., Germain, J., Barbas, S., et al. (2020). ABHD12 and LPCAT3 Interplay Regulates a Lyso-phosphatidylserine-C20:4 Phosphatidylserine Lipid Network Implicated in Neurological Disease. Biochemistry 59, 1793–1799. 10.1021/acs.biochem.0c00292.

22. Long, J.Z., and Cravatt, B.F. (2011). The metabolic serine hydrolases and their functions in mammalian physiology and disease. Chem Rev 111, 6022–6063.

23. Bachovchin, D.A., and Cravatt, B.F. (2012). The pharmacological landscape and therapeutic potential of the serine hydrolases. Nat Rev Drug Discov 11, 52–68.

24. Liu, Y., Patricelli, M.P., and Cravatt, B.F. (1999). Activity-based protein profiling: the serine hydrolases. Proc Natl Acad Sci U S A 96, 14694–14699.

25. Simon, G.M., and Cravatt, B.F. (2010). Activity-based proteomics of enzyme superfamilies: serine hydrolases as a case study. J Biol Chem. 285, 11051–11055.

26. Niphakis, M.J., and Cravatt, B.F. (2014). Enzyme inhibitor discovery by activity-based protein profiling. Annu Rev Biochem 83, 341–377. 10.1146/annurev-biochem-060713-035708.

27. Wu, C., Jin, X., Tsueng, G., Afrasiabi, C., and Su, A.I. (2016). BioGPS: building your own mash-up of gene annotations and expression profiles. Nucleic Acids Res 44, D313–316. 10.1093/nar/gkv1104.

28. Wu, C., Orozco, C., Boyer, J., Leglise, M., Goodale, J., Batalov, S., Hodge, C.L., Haase, J., Janes, J., Huss, J.W., 3rd, and Su, A.I. (2009). BioGPS: an extensible and customizable portal for querying and organizing gene annotation resources. Genome Biol 10, R130. 10.1186/gb-2009-10-11-r130.

29. Bachovchin, D.A., Ji, T., Li, W., Simon, G.M., Blankman, J.L., Adibekian, A., Hoover, H., Niessen, S., and Cravatt, B.F. (2010). Superfamily-wide portrait of serine hydrolase inhibition achieved by library-versus-library screening. Proc Natl Acad Sci U S A 107, 20941–20946. 10.1073/pnas.1011663107.

30. Kelkar, D.S., Ravikumar, G., Mehendale, N., Singh, S., Joshi, A., Sharma, A.K., Mhetre, A., Rajendran, A., Chakrapani, H., and Kamat, S.S. (2019). A chemical-genetic screen identifies ABHD12 as an oxidized-phosphatidylserine lipase. Nat Chem Biol 15, 169–178. 10.1038/s41589-018-0195-0.

31. Hoover, H.S., Blankman, J.L., Niessen, S., and Cravatt, B.F. (2008). Selectivity of inhibitors of endocannabinoid biosynthesis evaluated by activity-based protein profiling. Bioorg Med Chem Lett 18, 5838–5841. 10.1016/j.bmcl.2008.06.091.

32. Schlager, S., Vujic, N., Korbelius, M., Duta-Mare, M., Dorow, J., Leopold, C., Rainer, S., Wegscheider, M., Reicher, H., Ceglarek, U., et al. (2017). Lysosomal lipid hydrolysis provides substrates for lipid mediator synthesis in murine macrophages. Oncotarget 8, 40037–40051. 10.18632/oncotarget.16673.

33. Bradic, I., Kuentzel, K.B., Honeder, S., Grabner, G.F., Vujic, N., Zimmermann, R., Birner-Gruenberger, R., and Kratky, D. (2022). Off-target effects of the lysosomal acid lipase inhibitors Lalistat-1 and Lalistat-2 on neutral lipid hydrolases. Mol Metab 61, 101510. 10.1016/j.molmet.2022.101510.

34. Hamilton, J., Jones, I., Srivastava, R., and Galloway, P. (2012). A new method for the measurement of lysosomal acid lipase in dried blood spots using the inhibitor Lalistat 2. Clin Chim Acta 413, 1207–1210. 10.1016/j.cca.2012.03.019.

35. Hsu, K.L., Tsuboi, K., Adibekian, A., Pugh, H., Masuda, K., and Cravatt, B.F. (2012). DAGLbeta inhibition perturbs a lipid network involved in macrophage inflammatory responses. Nat Chem Biol. 8, 999–1007.

36. Hsu, K.L., Tsuboi, K., Chang, J.W., Whitby, L.R., Speers, A.E., Pugh, H., and Cravatt, B.F. (2013). Discovery and optimization of piperidyl-1,2,3-triazole ureas as potent, selective, and in vivo-active inhibitors of alpha/beta-hydrolase domain containing 6 (ABHD6). J Med Chem 56, 8270–8279. 10.1021/jm400899c.

37. Cognetta, A.B., 3rd, Niphakis, M.J., Lee, H.C., Martini, M.L., Hulce, J.J., and Cravatt, B.F. (2015). Selective N-Hydroxyhydantoin Carbamate Inhibitors of Mammalian Serine Hydrolases. Chem Biol 22, 928–937. 10.1016/j.chembiol.2015.05.018.

38. Blankman, J.L., Simon, G.M., and Cravatt, B.F. (2007). A comprehensive profile of brain enzymes that hydrolyze the endocannabinoid 2-arachidonoylglycerol. Chem Biol 14, 1347–1356. 10.1016/j.chembiol.2007.11.006.

39. Navia-Paldanius, D., Savinainen, J.R., and Laitinen, J.T. (2012). Biochemical and pharmacological characterization of human alpha/beta-hydrolase domain containing 6 (ABHD6) and 12 (ABHD12). J Lipid Res 53, 2413–2424. 10.1194/jlr.M030411.

40. Thomas, G., Betters, J.L., Lord, C.C., Brown, A.L., Marshall, S., Ferguson, D., Sawyer, J., Davis, M.A., Melchior, J.T., Blume, L.C., et al. (2013). The serine hydrolase ABHD6 Is a critical regulator of the metabolic syndrome. Cell Rep 5, 508–520. 10.1016/j.celrep.2013.08.047.

41. Cao, J.K., Kaplan, J., and Stella, N. (2019). ABHD6: Its Place in Endocannabinoid Signaling and Beyond. Trends Pharmacol Sci 40, 267–277. 10.1016/j.tips.2019.02.002.

42. Kano, K., Aoki, J., and Hla, T. (2022). Lysophospholipid Mediators in Health and Disease. Annu Rev Pathol 17, 459–483. 10.1146/annurev-pathol-050420-025929.

43. Yanagida, K., and Valentine, W.J. (2020). Druggable Lysophospholipid Signaling Pathways. Adv Exp Med Biol 1274, 137–176. 10.1007/978-3-030-50621-6_7.

44. Tan, S.T., Ramesh, T., Toh, X.R., and Nguyen, L.N. (2020). Emerging roles of lysophospholipids in health and disease. Prog Lipid Res 80, 101068. 10.1016/j.plipres.2020.101068.

45. Marrs, W.R., Blankman, J.L., Horne, E.A., Thomazeau, A., Lin, Y.H., Coy, J., Bodor, A.L., Muccioli, G.G., Hu, S.S., Woodruff, G., et al. (2010). The serine hydrolase ABHD6 controls the accumulation and efficacy of 2-AG at cannabinoid receptors. Nat Neurosci 13, 951–957. 10.1038/nn.2601.

46. Grabner, G.F., Fawzy, N., Pribasnig, M.A., Trieb, M., Taschler, U., Holzer, M., Schweiger, M., Wolinski, H., Kolb, D., Horvath, A., et al. (2019). Metabolic disease and ABHD6 alter the circulating bis(monoacylglycerol)phosphate profile in mice and humans. J Lipid Res 60, 1020–1031. 10.1194/jlr.M093351.

47. Rong, X., Wang, B., Dunham, M.M., Hedde, P.N., Wong, J.S., Gratton, E., Young, S.G., Ford, D.A., and Tontonoz, P. (2015). Lpcat3-dependent production of arachidonoyl phospholipids is a key determinant of triglyceride secretion. Elife 4. 10.7554/eLife.06557.

48. Hashidate-Yoshida, T., Harayama, T., Hishikawa, D., Morimoto, R., Hamano, F., Tokuoka, S.M., Eto, M., Tamura-Nakano, M., Yanobu-Takanashi, R., Mukumoto, Y., et al. (2015). Fatty acid remodeling by LPCAT3 enriches arachidonate in phospholipid membranes and regulates triglyceride transport. Elife 4, e06328. 10.7554/eLife.06328.

49. Adibekian, A., Martin, B.R., Chang, J.W., Hsu, K.L., Tsuboi, K., Bachovchin, D.A., Speers, A.E., Brown, S.J., Spicer, T., Fernandez-Vega, V., et al. (2012). Confirming target engagement for reversible inhibitors in vivo by kinetically tuned activity-based probes. J Am Chem Soc 134, 10345–10348. 10.1021/ja303400u.

50. Won, S.J., Davda, D., Labby, K.J., Hwang, S.Y., Pricer, R., Majmudar, J.D., Armacost, K.A., Rodriguez, L.A., Rodriguez, C.L., Chong, F.S., et al. (2016). Molecular Mechanism for Isoform-Selective Inhibition of Acyl Protein Thioesterases 1 and 2 (APT1 and APT2). ACS Chem Biol 11, 3374–3382. 10.1021/acschembio.6b00720.

51. Sugimoto, H., Hayashi, H., and Yamashita, S. (1996). Purification, cDNA cloning, and regulation of lysophospholipase from rat liver. J Biol Chem 271, 7705–7711. 10.1074/jbc.271.13.7705.

52. Wang, A., Deems, R.A., and Dennis, E.A. (1997). Cloning, expression, and catalytic mechanism of murine lysophospholipase I. J Biol Chem 272, 12723–12729. 10.1074/jbc.272.19.12723.

53. Sen, D., Maniyadath, B., Chowdhury, S., Kaur, A., Khatri, S., Chakraborty, A., Mehendale, N., Nadagouda, S., Sandra, U.S., Kamat, S.S., and Kolthur-Seetharam, U. (2023). Metabolic regulation of CTCF expression and chromatin association dictates starvation response in mice and flies. iScience 26, 107128. 10.1016/j.isci.2023.107128.

54. Chattopadhyay, T., Maniyadath, B., Bagul, H.P., Chakraborty, A., Shukla, N., Budnar, S., Rajendran, A., Shukla, A., Kamat, S.S., and Kolthur-Seetharam, U. (2020). Spatiotemporal gating of SIRT1 functions by O-GlcNAcylation is essential for liver metabolic switching and prevents hyperglycemia. Proc Natl Acad Sci U S A 117, 6890–6900. 10.1073/pnas.1909943117.

55. Talwadekar, M., Khatri, S., Balaji, C., Chakraborty, A., Basak, N.P., Kamat, S.S., and Kolthur-Seetharam, U. (2024). Metabolic transitions regulate global protein fatty acylation. J Biol Chem 300, 105563. 10.1016/j.jbc.2023.105563.

56. Rajendran, A., Vaidya, K., Mendoza, J., Bridwell-Rabb, J., and Kamat, S.S. (2020). Functional Annotation of ABHD14B, an Orphan Serine Hydrolase Enzyme. Biochemistry 59, 183–196. 10.1021/acs.biochem.9b00703.

57. Kumar, K., Mhetre, A., Ratnaparkhi, G.S., and Kamat, S.S. (2021). A Superfamily-wide Activity Atlas of Serine Hydrolases in Drosophila melanogaster. Biochemistry 60, 1312–1324. 10.1021/acs.biochem.1c00171.

