## Supplementary Figures for "Identification of ABHD6 as a regulator of lysophosphatidylserines in the mammalian liver and kidneys"

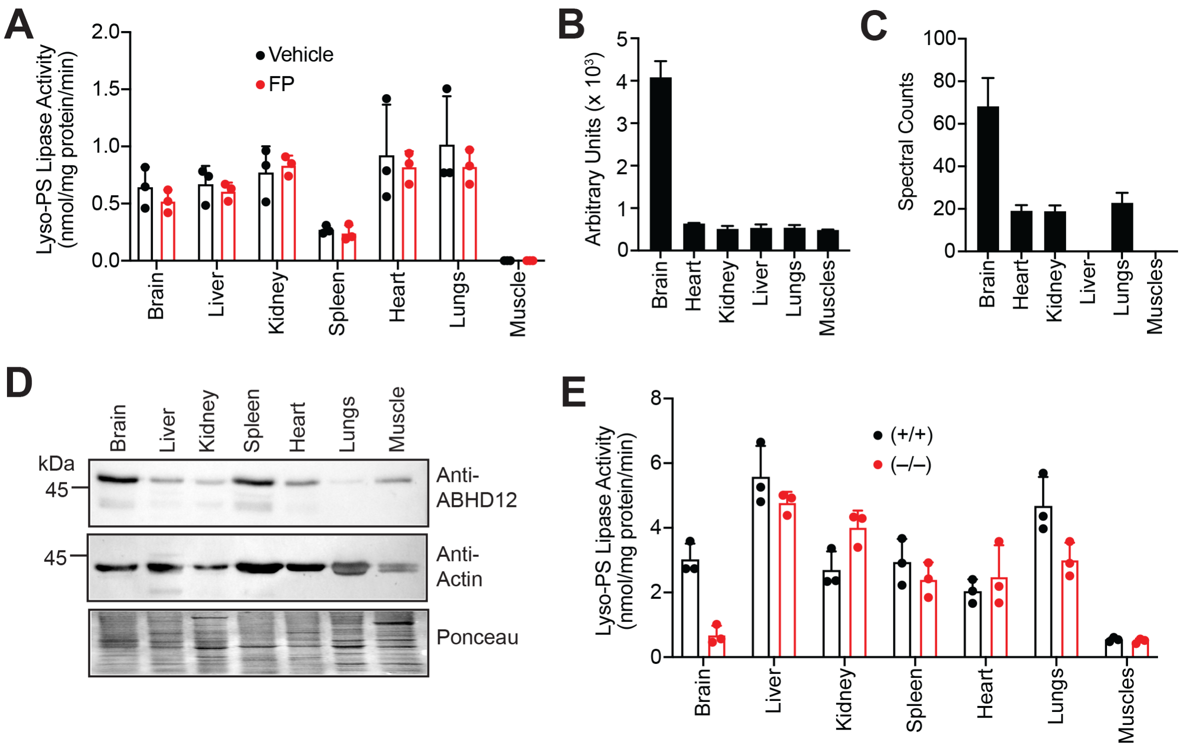


**Supplementary Figure 1. ABHD12 controls lyso-PS levels only in the mouse brain.** (**A**) The lyso-PS lipase activity in the soluble proteomic fractions of different mouse tissues treated with vehicle (DMSO) or FP-rhodamine (20 μM, 45 min). All assays were done using 20 μg of proteome against 100 μM C17:1 lyso-PS for 30 min at 37 °C. (**B**) The mRNA expression data exported from the large-scale gene expression database BioGPS[1,2], showing the heightened expression of ABHD12 in the mouse brain. (**C**) The activity of ABHD12 in various mouse tissues from a publicly available LC-MS based ABPP dataset (spectral counting-based data)[3], showing the heightened activity of ABHD12 in the membrane proteomic fraction of the mouse brain. (**D**) A representative western blot, showing the tissue distribution of ABHD12, further confirming highest protein levels on ABHD12 in the membrane proteomic fraction of the mouse brain consistent with data from (**B**) and (**C**). 50 μg of each membrane proteome was loaded onto the gel, and both β-Actin and Ponceau staining were used as loading controls for this experiment. This immunoblotting study was done three times with reproducible result each time. (**E**) The lyso-PS lipase activity in the membrane proteomic fractions of different mouse tissues obtained from wild type (+/+) or ABHD12 knockout (–/–) mice. All assays were done using 20 μg of proteome against 100 μM C17:1 lyso-PS for 30 min at 37 °C. All bar plots presented in this figure represents mean ± standard deviation from two (for **B**) or three (**A, C** and **E**) biological replicates.

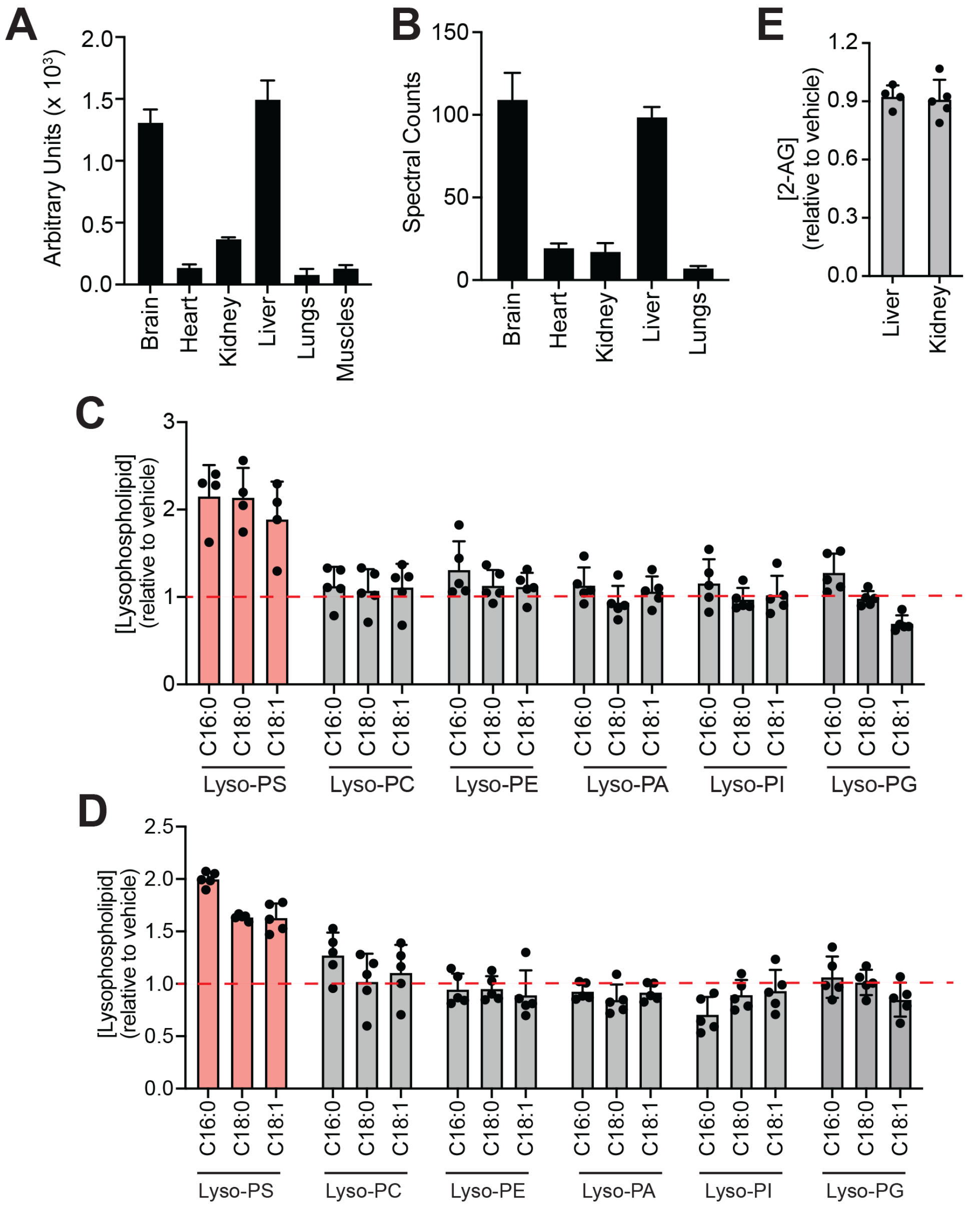


**Supplementary Figure 2. ABHD6 is a major lyso-PS lipase in the liver and kidney**. (A) The mRNA expression data exported from the large-scale gene expression database BioGPS[1,2], showing the heightened expression of ABHD6 in the mouse brain and liver, and to a lesser extent in the kidney. (**C**) The activity of ABHD6 in various mouse tissues from a publicly available LC-MS based ABPP dataset (spectral counting-based data)[3], showing the heightened activity of ABHD12 in the membrane proteomic fraction of the mouse brain and liver. (**C, D, E**) Relative levels of different lysophospholipids in the liver (**C**), and kidney (**D**) and 2-AG (**E**), upon KT185 treatment (10 mg/kg body weight, 4 hours), showing an increase of only lyso-PS, but no other lysophospholipid, from the selective inhibition of ABHD6. All bar plots presented in this figure represents mean ± standard deviation from two (for **A**) or four or more (**C, D** and **E**) biological replicates

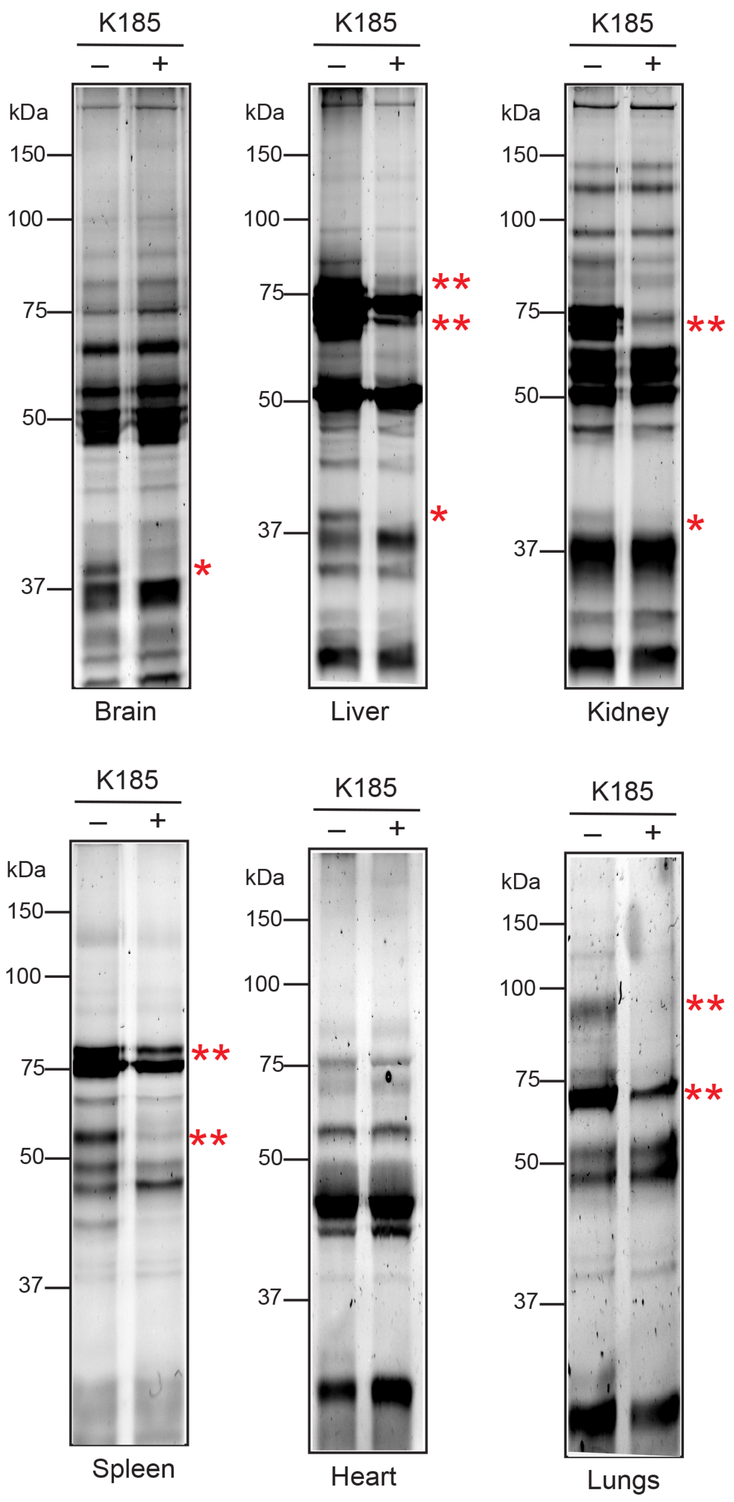


**Supplementary Figure 3.** Complete ABPP gels from **Figure 3A**, showing the loss of ABHD6 activity (band corresponding to the single red asterisk) in the brain, liver and kidney upon treatment with KT185 (orally, 10 mg/kg body weight, 4 hours). No ABHD6 activity was detected in the spleen, heart and lungs. The off-targets of KT185 (bands corresponding to two red asterisk) seen in the liver, kidney, spleen and lungs are likely enzymes from the CES-family (CES2 and CES3).

**REFERENCES.**


1. Wu, C., Jin, X., Tsueng, G., Afrasiabi, C., and Su, A.I. (2016). BioGPS: building your own mash-up of gene annotations and expression profiles. Nucleic Acids Res *44*, D313-316. 10.1093/nar/gkv1104.

2. Wu, C., Orozco, C., Boyer, J., Leglise, M., Goodale, J., Batalov, S., Hodge, C.L., Haase, J., Janes, J., Huss, J.W., 3rd, and Su, A.I. (2009). BioGPS: an extensible and customizable portal for querying and organizing gene annotation resources. Genome Biol *10*, R130. 10.1186/gb-2009-10-11-r130.

3. Bachovchin, D.A., Ji, T., Li, W., Simon, G.M., Blankman, J.L., Adibekian, A., Hoover, H., Niessen, S., and Cravatt, B.F. (2010). Superfamily-wide portrait of serine hydrolase inhibition achieved by library-versus-library screening. Proc Natl Acad Sci U S A *107*, 20941-20946. 10.1073/pnas.1011663107.
